# Systemic evidence of acute seizure-associated neuronal injury in patients with temporal lobe epilepsy

**DOI:** 10.1101/184366

**Authors:** Shailaja Kunda, Reghann G LaFrance-Corey, Fatemeh Khadjevand, Gregory A Worrell, Charles L Howe

## Abstract

Patients with drug refractory temporal lobe epilepsy frequently accumulate cognitive impairment over time, suggesting neuronal loss induced by seizures. We measured serum levels of neuron-specific enolase (NSE), a neuronal injury marker, relative to levels of S100β, a marker of glial injury, at 6 AM, 9 AM, noon, 3 PM, and 6 PM over the course of several days in 7 epilepsy patients and 4 healthy controls. All epilepsy patients exhibited significant deviations in NSE levels through time, and 4 of the epilepsy patients exhibited large sample entropy values and large signal variation metrics for NSE relative to S100β. Controls did not exhibit such changes. Correlation analysis revealed that NSE levels were significantly elevated after clinical seizure events. There was also a highly significant relationship between increased EEG spike frequency and an increase in serum NSE levels measured 24 hours later. The detection of large but transient post-ictal increases in NSE suggests that even self-limited seizures may cause an injury to neurons that underlies cognitive decline in some patients. Post-ictal assessment of serum NSE may serve as a biomarker for measuring the efficacy of future acute neuroprotective strategies in epilepsy patients.

## Introduction

More than 30% of all patients with epilepsy continue to experience seizures despite treatment with a wide range of anti-epileptic drugs^1^. In these refractory patients, a subset exhibit a progressive disease phenotype, both with regard to increasing seizure frequency over time and from the perspective of accumulating cognitive impairment^2; 3^. Indeed, epilepsy for some patients is effectively a neurodegenerative disorder^4^. This is particularly true in patients with temporal lobe epilepsy marked by mesial temporal sclerosis^5^, and several studies indicate that progressive hippocampal atrophy as assessed by MRI correlates with increasing seizure frequency and cognitive decline in these patients^6-10^. In experimental models of epilepsy, induction of status epilepticus, not surprisingly, leads to hippocampal neuron loss^11^. However, spontaneously recurring seizures in such models are also associated with neuronal loss^12^, suggesting that individual seizures may induce neurodegeneration. In humans, neuronal injury induced by trauma, hypoxia, and stroke can be detected by measuring levels of neuron-specific enolase (NSE) in serum^13^. Building on previous work assessing NSE levels following seizures^14-17^, in this study we collected serial blood samples from epilepsy patients and healthy control subjects and measured changes in both NSE and the glial injury marker S100β^13^ through time in an effort to correlate seizures and electroencephalographic events with neuronal injury.

## Materials and Methods

### Subjects and study design

Study protocols were approved by the Mayo Clinic institutional review board and all experiments were performed in accordance with the relevant guidelines and regulations. All subjects provided written informed consent. Patients with intractable focal epilepsy were admitted to the Mayo Clinic epilepsy monitoring unit (EMU) for routine diagnostic computer-assisted continuous video-EEG recording. Control subjects were admitted to the Mayo Clinic clinical research unit (CRU). Subjects in both groups were between 18 and 65 years old. Individuals were excluded on the basis of pregnancy, weight less than 110 lbs, history of chronic illness (other than epilepsy), active malignancy, active infection, or history of immunosuppressive therapy within 6 months of the study. Control subjects were further excluded on the basis of seizure history. Patients and controls received a peripheral venous catheter at the start of the study and a blood sample was collected immediately for complete blood count and differential. For all subjects, blood was collected at 6 AM, 9 AM, noon, 3 PM, and 6 PM throughout the study duration. Contingencies included delaying sample collection by 30 minutes during an active clinical seizure at the normal draw time, up to two replacement intravenous lines during the study, and conversion to venipuncture upon repeated intravenous line failure. Samples were collected into rapid serum separator tubes (BD 368774), immediately inverted 6 times, then transported to the research lab at room temperature. Within 30 minutes of collection the samples were centrifuged and the serum fraction was aliquoted and stored at −80ºC.

### Serum analysis

Frozen samples were thawed on ice and clarified by high speed centrifugation (10000 x g, 5 min). Once thawed, sample aliquots were never refrozen or reused. All samples were visually inspected for hemolysis (none exhibited obvious signs); a subset of samples were analyzed for hemolysis using the method of Harboe^18^. Briefly, serum was diluted 11-fold in PBS and hemoglobin was measured based on the following equation: C_HB_ = 1.65(A_415_) − 0.93(A_380_) − 0.73(A_450_). Samples exhibited 0.03 ± 0.02 mg/mL hemoglobin (n=25), which is within the normal range (0.02 ± 0.02 mg/mL)^18^. Levels of NSE (Alpco 43-NSEHU-E01) and S100ß (Millipore EZHS100B-33K) were determined by enzyme-linked immunosorbent assay following manufacturer’s directions. Standard curves were analyzed for all assays. Across all analyses the intra-and inter-assay coefficient of variation was less than 15%.

### Seizure and spike frequency analysis

Continuous video-EEG was collected over multiple days using 32 scalp electrodes (modified 10-20 montage; 250 Hz sampling rate) (Natus Medical Inc). Differential amplifiers with band-pass filters between 1 and 70 Hz were used to minimize the effects of high frequency and low frequency artifacts. A vertex recording reference and ground was used during acquisition. Clinical seizure events were identified by visual inspection of the EEG coupled to video analysis. Interictal epileptiform discharges were assessed by visual review in referential, bipolar, and Laplacian montages using digital reformatting of the EEG. For automated analyses the archived EEG files from the EMU were pre-processed in Natus Xltek software and the individual files were aligned by timestamp to permit association with the serum measurements. The continuous spike frequency was quantified using the automated spike detection algorithm available in Persyst 13 (www.persyst.com). This algorithm uses approximately 20 feedforward neural network rules to characterize relevant events on a common electrode referential montage and, in parallel, on montages referenced to (Fp1+Fp2), (T3+T4), and (O1+O2). A detailed methodology and performance assessment for this spike detector was recently published^19^.

### Analysis of sample entropy and relative signal variation

To assess the significance of dynamic changes in the level of NSE measured in patient samples we calculated sample entropy following the protocol of Richman and Moorman^20^. This method, which reveals “novelty” in time series data, is based on the conditional probability that two vector sequences derived from the same time series will be roughly similar, to within some predefined tolerance. Sample entropy, derived from the original concept of approximate entropy^21^, provides an entropy measure for relatively short and noisy biological time series data. Following the guidelines established by Yentes and colleagues^22^, we determined the optimal tolerance parameter for the very short time series data collected in our patient and healthy control cohorts. The very short series in our study did tend to display extreme behavior under certain parameter constraints (blowing up to infinity, for example), but the use of an iterative modeling process using random and patterned sequences provided a working algorithm. In brief, using a script written in Matlab, each time series was parsed into an array of overlapping vectors comprised of 2 and 3 sequential points. The Chebyshev distance between each vector in the array was calculated and compared to a tolerance factor, r, that was empirically established as 0.6 times the standard deviation of all the experimental measurements in the study (NSE or S100β). Explicitly following the procedure of Richman and Moorman, the probability of vector similarity at length 2 and length 3 was calculated and the sample entropy was taken as the negative natural logarithm of the ratio of the 3-length probability to the 2-length probability^20^. The algorithm was validated on sequences of 5000 random numbers drawn from a normal distribution with the mean and standard deviation of the experimental samples, with the average sample entropy of 1000 iterations of this calculation matching the Richman and Moorman values^20^. In addition, because the sample entropy was sensitive to time series length, we established a normalization factor for series of lengths between 5 and 13 values using signals with no entropy (sequences of the same repeated number). Because the absolute sample entropy values are lacking in context, we also calculated a relative signal variation metric (SVM) by taking the ratio of the NSE sample entropy (SE_NSE_) to the S100β (SE_S100β_) sample entropy measured in the same patient:

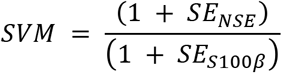

To prevent division by zero (when the S100β series displayed essentially no variation), all sample entropies were scaled such that no variation (low entropy) was equivalent to one.

### Analysis of changes in NSE levels and spike frequency

Serum NSE levels were recast as the change in concentration relative to the preceding NSE measurement. This delta was then recast as the number of standard deviations in NSE derived from the 4 CRU subjects. This value was binarized so that NSE changes greater than or equal to 3 standard deviations were set to 1 and all other values were set to zero. The absolute number of EEG spikes was organized into one hour epochs preceding each NSE measurement. This frequency value was binarized so that any frequency greater than 10 spikes/hr was set to 1 and all other values were set to zero. Missing values (due to the absence of sufficient EEG collection time prior to the first several NSE measurements) were maintained as empty cells. The relationship beween these binarized values was assessed using an estimated maximum likelihood logistic model on a binomial distribution to generate the *χ*^2^ significance values shown in Figure 5E. A standard least squares linear regression model was used to determine R^2^ and measure the analysis of variance; this model was also used to visualize the associations shown in Figure 5D. Power was determined from the leverage plot.

### Statistics

Curran-Everett guidelines were followed^23^. Statistical analyses were performed using JMP Pro 12 (SAS Institute Inc). Post hoc power analysis was performed for all experiments. Normality was determined by the Shapiro-Wilk test and normally distributed data were checked for equal variance. Parametric tests were only applied to data that were both normally distributed and of equal variance. NSE measurements in the EMU and CRU samples were analyzed by one-way ANOVA using Dunnett’s pairwise comparison to the aggregated CRU values (Bonferroni adjusted P-value). Correlations between NSE and S100β deviations from median, NSE vs S100β levels through time, and NSE vs seizure time were performed using least squares linear regression modeling with effect leveraging and analysis of variance. Correlations for NSE vs spike epoch were generated using a generalized logistic model on a binomial distribution. The single variable reduction that resulted from calculation of the signal variation metric was analyzed by t-test (data normally distributed). Ranges in all graphs reflect the 95% confidence interval. Where reported, all R^2^ values are adjusted for sample size.

## Results

### Study subject characteristics

Between 2013 and 2016 seven patients admitted to the Mayo Clinic epilepsy monitoring unit (EMU) for continuous video-electroencephalography (EEG) monitoring as part of standard clinical care for intractable focal epilepsy were enrolled in a research study to longitudinally collect serum samples for analysis of systemic neural injury markers (Table 1). Inclusion in the subsequent analysis required evidence of at least one clinical seizure during the study. Subjects ranged in age from 25 to 49 years, were evenly distributed by sex, and had disease durations that ranged from 5 months to 41 years. Of the 7 subjects, 5 had clear evidence of mesial temporal sclerosis. Between 2016 and 2017 four control subjects were enrolled at the Mayo Clinic clinical research unit (CRU) for longitudinal collection of serum samples to compare to the EMU subjects (Table 2). Subjects ranged in age from 19 to 61 years (3 females, 1 male) and had no history of seizures or epilepsy; other chronic disease conditions were not screened. In addition to the longitudinally sampled control subjects, 41 non-neurological control serum samples were acquired from the Mayo Clinic Center for Individualized Medicine Biobank biorepository. These controls (HC) ranged in age from 19 to 75 (34 females, 7 males).

**Table 1.**
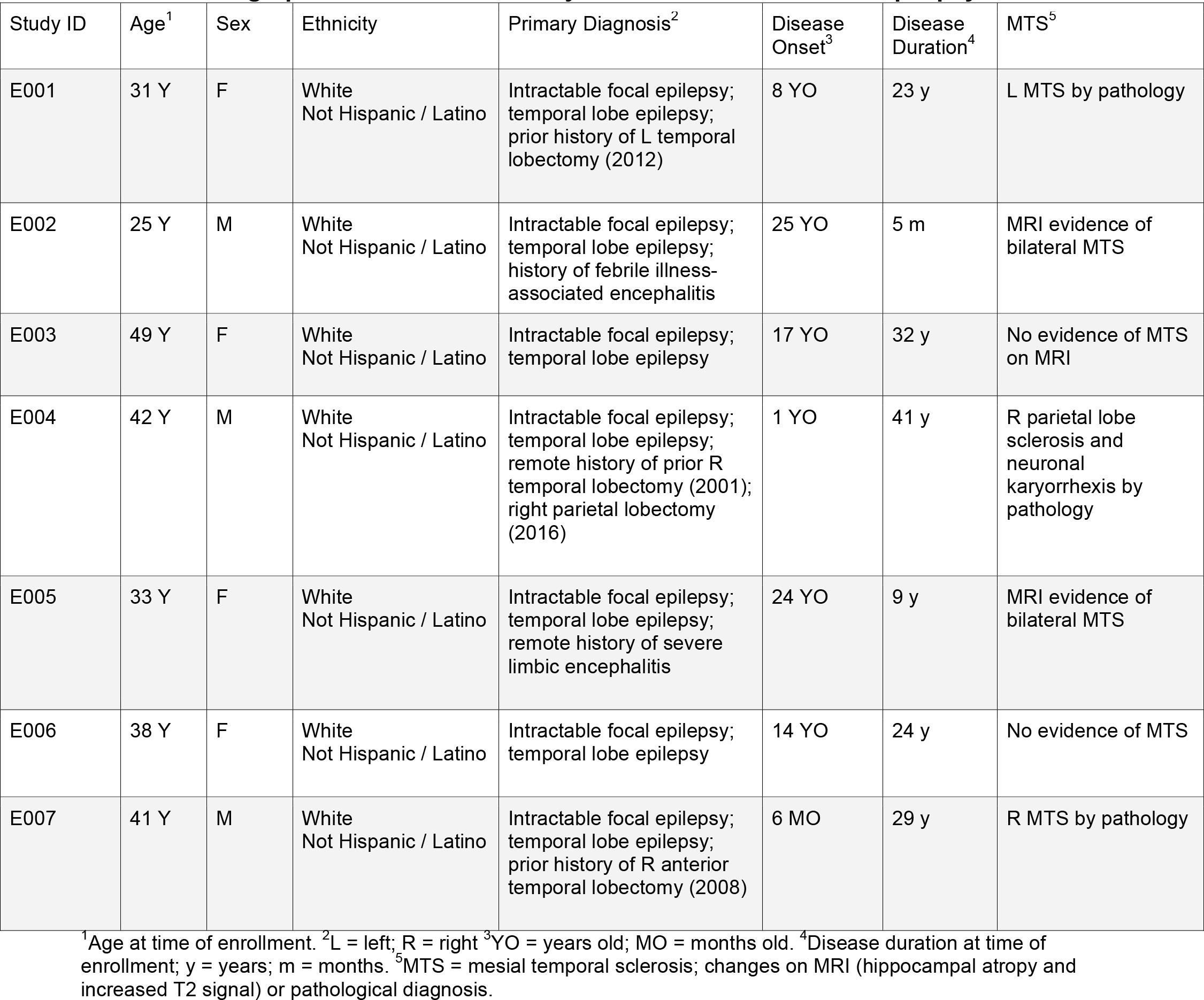
**Demographic information for subjects with intractable focal epilepsy**

| Study ID | Age<sup>1</sup> | Sex | Ethnicity | Primary Diagnosis<sup>2</sup> | Disease Onset<sup>3</sup> | Disease Duration<sup>4</sup> | MTS<sup>5</sup> |
| --- | --- | --- | --- | --- | --- | --- | --- |
| E001 | 31 Y | F | White<br>Not Hispanic / Latino | Intractable focal epilepsy;<br>temporal lobe epilepsy;<br>prior history of L temporal lobectomy (2012) | 8 YO | 23 y | L MTS by pathology |
| E002 | 25 Y | M | White<br>Not Hispanic / Latino | Intractable focal epilepsy;<br>temporal lobe epilepsy;<br>history of febrile illness-associated encephalitis | 25 YO | 5 m | MRI evidence of bilateral MTS |
| E003 | 49 Y | F | White<br>Not Hispanic / Latino | Intractable focal epilepsy;<br>temporal lobe epilepsy | 17 YO | 32 y | No evidence of MTS on MRI |
| E004 | 42 Y | M | White<br>Not Hispanic / Latino | Intractable focal epilepsy;<br>temporal lobe epilepsy;<br>remote history of prior R temporal lobectomy (2001);<br>right parietal lobectomy (2016) | 1 YO | 41 y | R parietal lobe sclerosis and neuronal karyorrhexis by pathology |
| E005 | 33 Y | F | White<br>Not Hispanic / Latino | Intractable focal epilepsy;<br>temporal lobe epilepsy;<br>remote history of severe limbic encephalitis | 24 YO | 9 y | MRI evidence of bilateral MTS |
| E006 | 38 Y | F | White<br>Not Hispanic / Latino | Intractable focal epilepsy;<br>temporal lobe epilepsy | 14 YO | 24 y | No evidence of MTS |
| E007 | 41 Y | M | White<br>Not Hispanic / Latino | Intractable focal epilepsy;<br>temporal lobe epilepsy;<br>prior history of R anterior temporal lobectomy (2008) | 6 MO | 29 y | R MTS by pathology |

<sup>1</sup>Age at time of enrollment. <sup>2</sup>L = left; R = right <sup>3</sup>YO = years old; MO = months old. <sup>4</sup>Disease duration at time of enrollment; y = years; m = months. <sup>5</sup>MTS = mesial temporal sclerosis; changes on MRI (hippocampal atrophy and increased T2 signal) or pathological diagnosis.

**Table 2.** **Control subject demographic information**

| Study ID | Age* | Sex | Ethnicity | Primary Diagnosis |
| --- | --- | --- | --- | --- |
| C001 | 19 Y | F | White<br>Not Hispanic/ Latino | Abnormal uterine bleeding |
| C002 | 40 Y | M | White<br>Not Hispanic/ Latino | Head Tremor (Essential Tremor)<br>Remote history of visual loss (migraine equivalent for visual loss) |
| C003 | 61Y | F | White<br>Not Hispanic/ Latino | Conjunctivitis, OS likely related to dry eyes |
| C004 | 53 Y | F | White<br>Not Hispanic/ Latino | Asthma |

\*Age at time of study enrollment.

### NSE and S100β levels in epilepsy patients versus healthy controls

NSE and S100β were measured by ELISA in clarified serum samples. The distributions of both factors failed normality testing (NSE, Shapiro-Wilks W=0.6415, P<0.0001; S100β, Shapiro-Wilks W=0.9645, P=0.0023) so only non-parametric statistical tests were applied. The CRU controls, grouped irrespective of collection time, had 17.1 ± 1.3 ng/mL [13.3, 22.0] NSE and 66.3 ± 6.9 pg/mL [45.5, 90.3] S100β. Figure 1 shows the mean ± 95% CI for NSE (Figure 1A) and S100β (Figure 1B) in the grouped CRU samples (light blue band) overlaid with each individual measurement in the control and experimental groups. Statistical analysis of the NSE measurements (across all draws irrespective of time) revealed that only EMU subject E005 was significantly different from the grouped CRU controls (F=4.3228, P=0.0004 by one-way ANOVA; E005 vs CRU at P=0.0024 by Dunnetťs pairwise method vs control; power = 0.986). The absence of significant differences in the epilepsy patients as compared to the controls, despite more dispersion in the NSE measurements in the EMU subjects, suggests that analysis of NSE levels in the absence of consideration for temporality is insufficient to discriminate patients with epilepsy from healthy controls.

**Figure 1.**
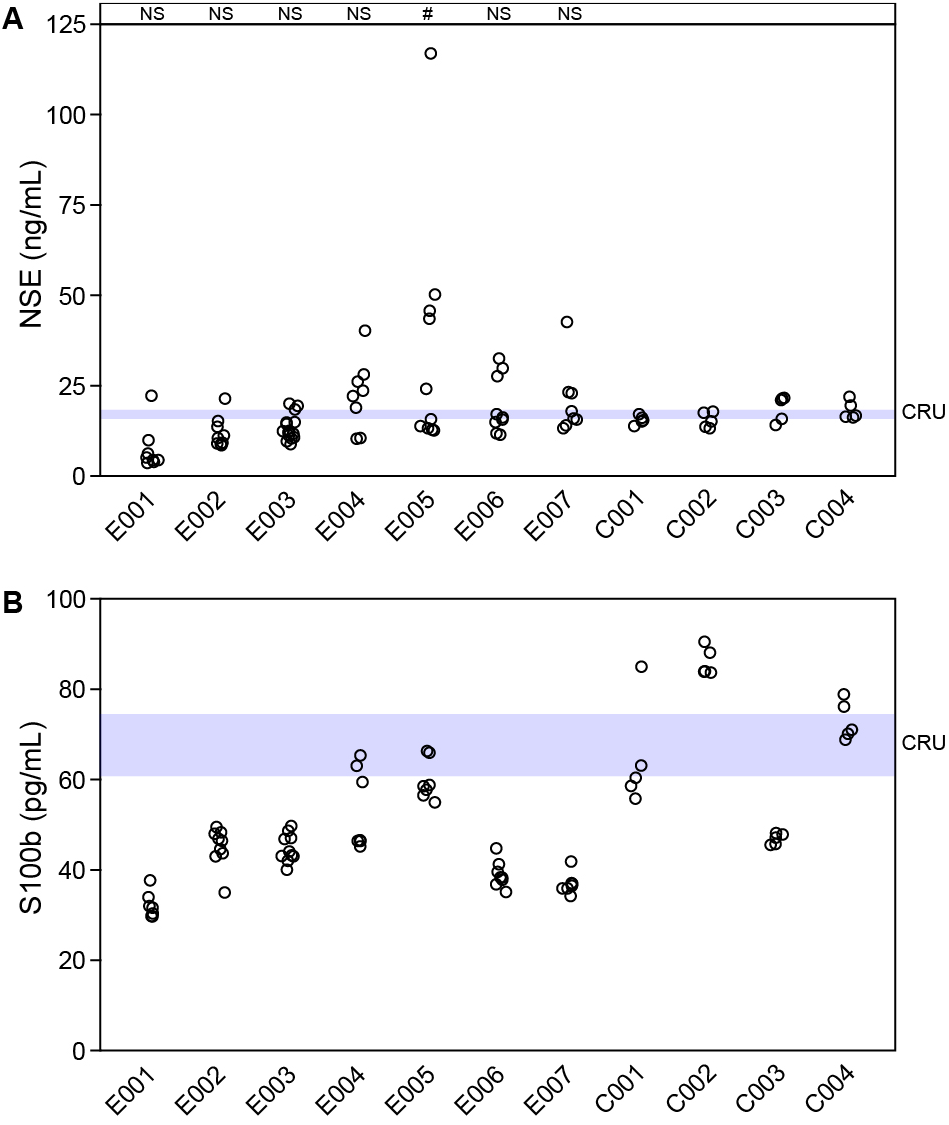
Serum levels of NSE and S100β in epilepsy patients and healthy controls are not different when analyzed in aggregate. A) Multiple serum samples were collected from 7 patients in the epilepsy monitoring unit (EMU) at different times (6 AM, 9 AM, noon, 3 PM, and 6 PM) during several days of monitoring; each sample is represented by one dot. NSE was measured in serum collected from 4 healthy control subjects in the clinical research unit (CRU) at 6 AM, 9 AM, noon, 3 PM, and 6 PM during one day to establish a reference range for samples collected under conditions identical to the EMU patients (blue bar shows mean ± 95% Cl; each sample is represented by one dot). Data are not normally distributed (W=0.6415, P<0.001 by Shapiro-Wilk test). One-way ANOVA with Dunnetťs pairwise comparison to group CRU controls revealed that NSE levels were only signficantly elevated in 1 of the 7 EMU patients (# = P<0.01; NS = not signficant). B) The same EMU and CRU serum samples used for NSE were assessed for S100β (blue bar shows mean ± 95% Cl for CRU controls). S100β levels were not elevated in any of the EMU patients and were, in fact, relatively lower in some patients.

### Temporal changes in NSE are not correlated with S100β levels

The primary goal of this study was to measure serum markers of neural injury repeatedly in the same subject over time in order to capture transient changes associated with acute seizure-induced injury. Figure 2 shows that all 7 EMU patients exhibited apparent “spikes” in NSE detected within serum over the course of several days (Figure 2A-2G; note the extended y-axis scale in 2E). At the same time points, S100β levels in the same subject were relatively stable. In contrast, the levels of both NSE and S100β measured in the CRU control subjects over the course of one day remained relatively stable (Figure 2H). The pattern of NSE levels in the CRU subjects suggested that diurnal rhythmicity did not explain the transient changes observed in the EMU patients. However, to verify that the changes in NSE levels were not tied to a daily cycle, the absolute level of NSE measured at each timepoint was normalized to the maximum NSE level measured across all timepoints to give an intrasubject percent of maximum value. Plotting these relative levels across time revealed no apparent cyclic pattern of maxima or minima in the NSE levels (Figure 3A). Likewise, given the age range in the study, the amount of NSE (Figure 3B) and S100β (Figure 3C) for each single draw healthy control (HC) subject was plotted against age and a line was fit by regression analysis. Neither factor exhibited an age-dependency, though a very weak trend between increasing serum S100β and age may be present in the control population. Finally, to verify that the response profile observed in the EMU patients was not the result of age, the median amount of NSE (Figure 3D) and S100β (Figure 3E) measured for each subject was plotted against age at the time of collection. No obvious trends were observed (NSE: R^2^=0.05; S100β: R^2^<0.0001), suggesting that the transient spikes in serum NSE observed in the EMU cohort were not a factor of either time-of-day or subject age.

**Figure 2.**
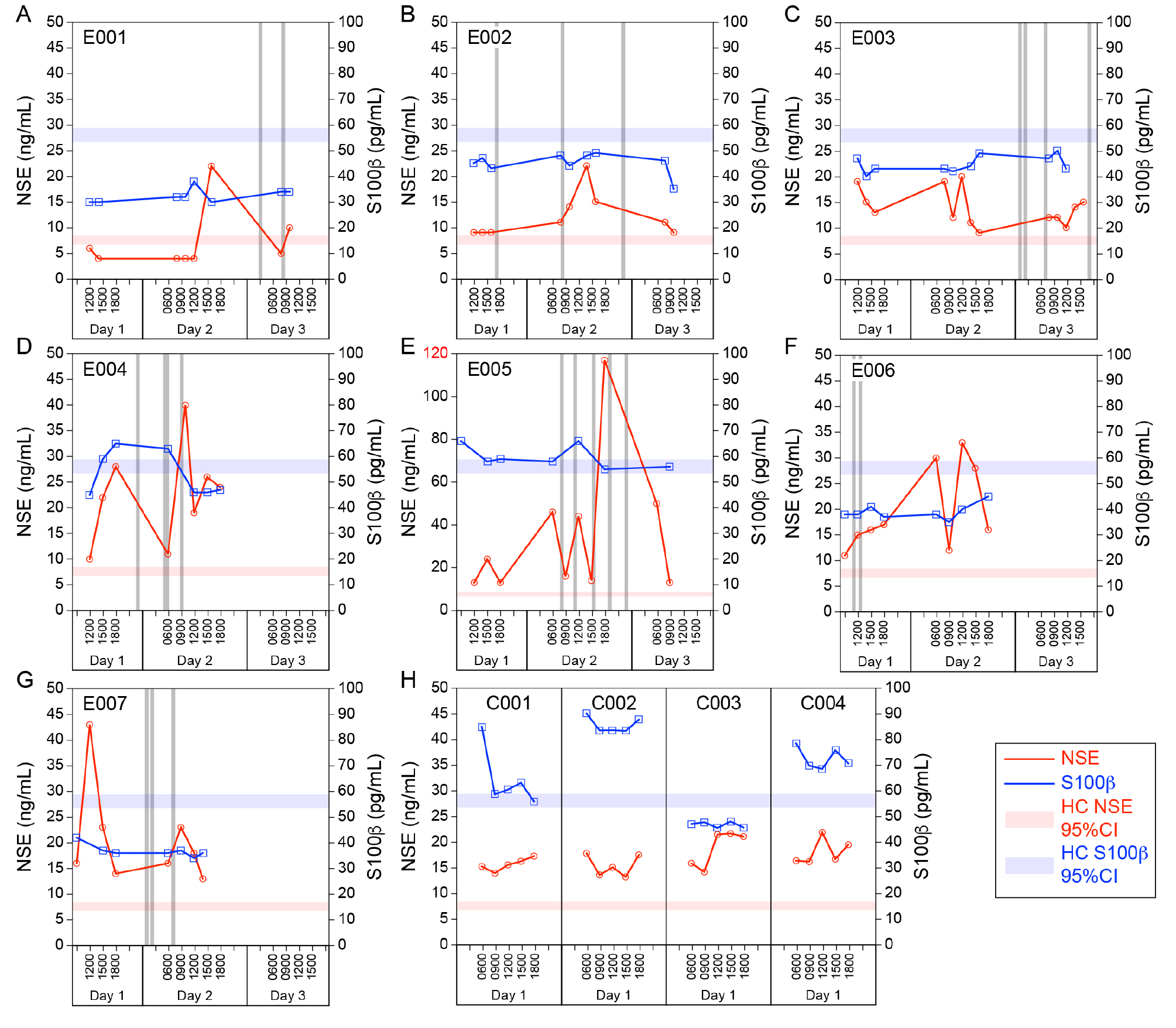
“Spikes” in serum NSE levels are observed in epilepsy patients but not in healthy controls or in S100β levels. Serially drawn blood samples from epilepsy patients (E001-E007, panels A-G) and healthy controls (C001-C004, panel H) were used to establish individual patterns of NSE (red lines) and S100β (blue lines) in serum through time. Sampling times were restricted to 0600, 0900, 1200, 1500, and 1800 hours; for EMU patients the draws continued throughout the duration of EEG monitoring. All panels are scaled to 50 ng/mL NSE (left axis) and 100 pg/mL S100β (right axis), except for E005 (E; 120 ng/mL NSE). The horizontal light red bars in all panels represent mean± 95% Cl for NSE in single-draw healthy controls (HC); the horizontal light blue bars in all panels represent mean± 95% Cl for S100β in single-draw healthy controls (HC). Vertical gray bars represent clinical seizure events.

**Figure 3.**
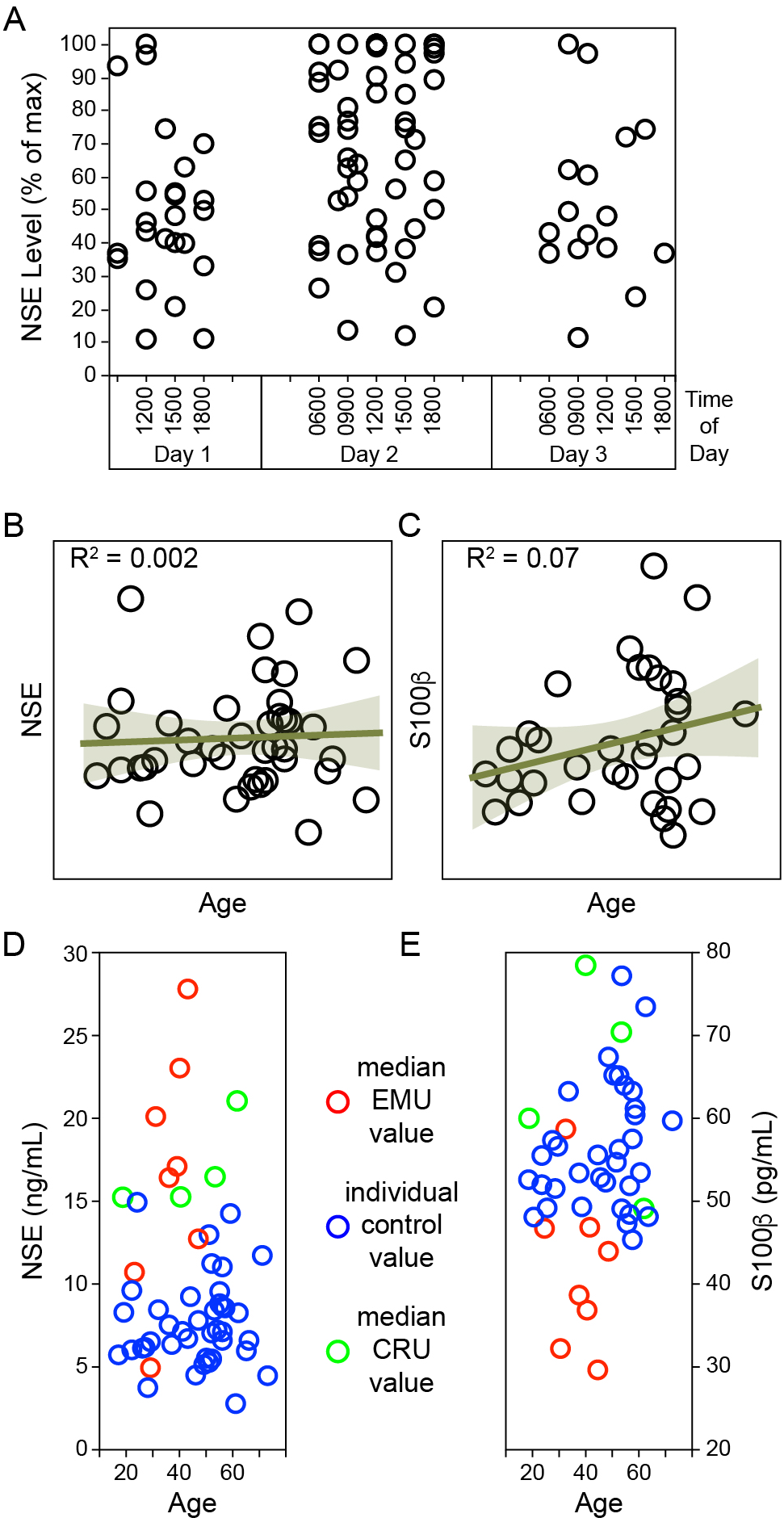
NSE and S100β levels are not associated with time-of-day or age of subject. A) Each NSE measurement for all EMU and CRU subjects was converted to percent of maximum for the individual and plotted against time-of-day (CRU samples are shown on day 2 in order to capture the entire 6 AM to 6 PM scale). There is no apparent pattern in the distribution of maximum or minimum NSE levels, suggesting that time-of-day did not drive the variations observed in the EMU patients. B) Absolute NSE values measured in 40 single-draw healthy controls were plotted against age at time of blood collection, revealing no relationship (R^2^=0.002). C) S100β levels in 34 single-draw healthy control subjects were also not correlated with age (R^2^=0.07). D) The median NSE level for each individual EMU (red) and CRU (green) subject was also plotted against age and overlaid with the single sample healthy control values (blue). Again, no apparent relationship between age and serum NSE was revealed. E) Similarly, no relationship between median serum S100β and age was apparent in EMU or CRU subjects.

In order to evaluate the significance of the temporal changes measured in the EMU patients, we employed four strategies to determine whether the variation in NSE levels was independent of and greater than that observed in S100β. In the first, each of the measurements for NSE and S100β in the EMU subjects were converted to the absolute value of the deviation from the median across all measurements within the same subject. An ANOVA was then performed using a standard least squares fit with patient and serum analyte as model effects. This analysis revealed that there was a significant effect in the cohort (F=3.3184, P=0.0017) and that the deviations in NSE were significantly larger than the S100β deviations (P=0.0064 by t-test). In the second, the linear dependence between NSE and S100β time series was assessed for each EMU subject by calculating the correlation coefficient. None of the patients exhibited a significant correlation between serum analytes (R range [−0.3708, 0.4594], P range [0.2135, 0.9060]), indicating that the changes in NSE levels were not associated with similar changes in S100β levels. However, only one of the CRU subjects showed a significant correlation between NSE and S100β (C002, R=0.9529, P=0.024), suggesting that this method is not sufficiently sensitive to robustly rule out a relationship between the serum analytes. In the third strategy, the time series data were converted to percent of maximum value measured for each analyte in each patient. Centering the normalized curves on the maximum measurement for NSE (time 0) revealed a high degree of signal variation for this factor that is not observed in the S100β curves (Figure 4A). Building on this, in the fourth strategy we calculated the sample entropy for each time series in each patient and used these values to calculate a signal variation metric (Figure 4B). None of the EMU or CRU S100β time series exhibited high sample entropy values (taken as >0.5; though see C001). In contrast, E001, E004, E005, and E006 had large NSE sample entropies and these same subjects exhibited large signal variation metrics. Using the signal variation metric to reduce each factor in each patient to a single value revealed that the EMU subjects were significantly different from the CRU controls (Figure 4B; P=0.0004 by t-test; power = 0.905).

**Figure 4.**
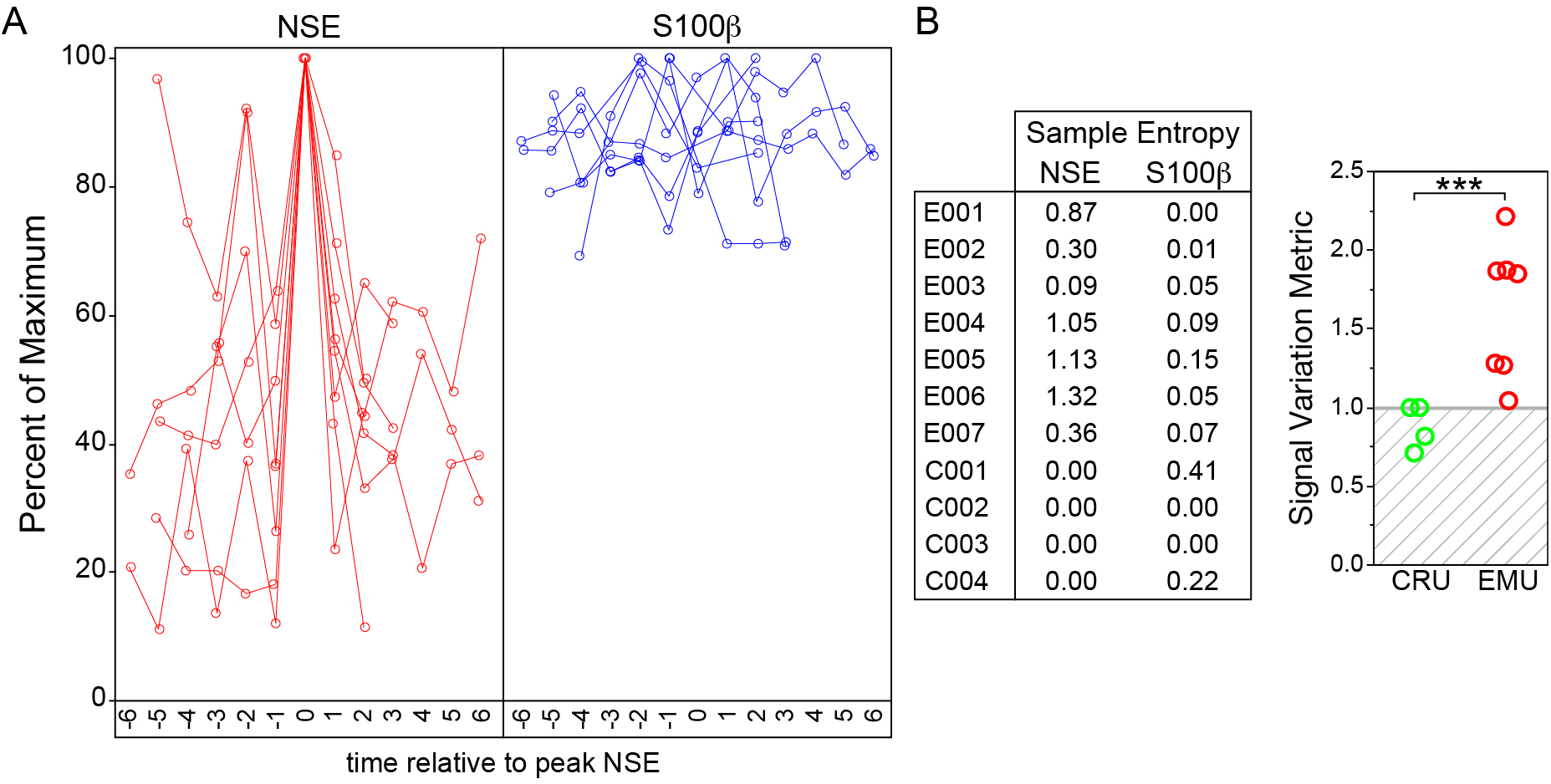
NSE levels exhibit high sample entropy and large signal variation in epilepsy patients but not healthy controls. A) NSE (red lines) and S100β (blue lines) measurements were converted to percent of maximum for each individual. The time at which the peak NSE value (100%) was measured in each subject was defined as t=0 and the remaining measurements were plotted relative to this timepoint. S100β measurements were aligned based on the t=0 set for NSE. While the NSE measurements exhibit a clear spike phenotype centered on t=0, the S100β values show no pattern, indicating that the high signal variability in NSE is not the result of non-specific serum changes. B) Sample entropy was calculated for NSE and S100β measurements in each subject. Most of the EMU patients exhibited high sample entropy (>0.5) while all of the CRU subjects had zero entropy in the NSE measurements. All S100β measurements showed low sample entropy. To further reduce the measurements to a single metric, the signal variation was calculated for each individual. Signal variations less than or equal to 1.0 indicate either no variability in the NSE measurements or variability present in both the NSE and S100β values. All of the CRU subjects had signal variation metrics below 1.0; all of the EMU patients had signal variations above 1.0, with E001, E004, E005, and E006 showing high signal variation.

### Changes in NSE level are temporally associated with clinical seizures and with electroencephalographic spiking

To characterize the relationship between clinical seizure events and serum NSE levels, the absolute NSE concentrations were converted to percent of the maximum measured for each patient and these values were temporally realigned to the first, second, or third seizure event determined by video scalp EEG (Figure 5A-5C). In effect, the first seizure was set to time zero for each patient and all of the NSE measurements were plotted relative to this time (time before seizure and time after seizure). The distribution of NSE against relative seizure time was then analyzed by least squares linear regression. Relative to the first seizure, a positive correlation (R^2^=0.143) was observed between time after seizure and increased NSE levels (Figure 5A). This effect was significant at P=0.0064 by ANOVA (F=8.0948; power = 0.797) and at P<0.0001 by *χ*^2^ analysis. Likewise, relative to the second seizure, a positive correlation (R^2^=0.127; P=0.0105; F=7.0613; power = 0.741) was still observed between time after seizure and elevated NSE (Figure 5B). However, by the third seizure event no correlation was detected (R^2^=0.015; P=0.5238; F=0.3855; power = 0.093; Figure 5C). This suggests that, in general, NSE levels increased after the first or second clinical seizure event.

While changes in NSE levels were temporally correlated with preceding overt seizures, there were also NSE spikes that exhibited an apparent disconnection from clinical events. Moreover, the limited number of clinical seizure events prevented fine resolution analysis of the time from seizure to NSE changes. In order to determine if electroencephalographic events that did not necessarily manifest as seizures were also related to NSE changes we measured the continuous spike frequency in the EEG from four EMU patients (E004-E007; data were not available for E001, E002, E003). For this analysis the data were reduced as described in the methods to yield a unitless measure of increased NSE level (relative to the preceding measurement) and a unitless measure of increased spike frequency. The spike frequency values were binned into one hour epochs relative to the NSE measurements (e.g. 0-1 hr before, 1-2 hr before). Due to the length of the recordings available for the patients the longest time used for analysis was 30 hr before each NSE measurement. The relationship between NSE changes and spike frequency was characterized using an estimated maximum likelihood model on a binomial distribution and by least squares linear regression. The regression fits for every epoch from −1 hr to −30 hr relative to the NSE measurement revealed a strong association between increased serum NSE and spiking on the EEG 24 hr earlier (R^2^=0.595; Figure 5D). Associations were also detected at 23 hr before the serum measurement and at −18 and −15 hr (Figure 5D). The 24 hr association was highly significant by ANOVA (F=22.056; P=0.0003; power = 0.992; Figure 5E). The other associations were significant at P<0.05 but were underpowered. These findings suggest that a period of spiking activity results in elevated serum NSE levels after a delay of about 24 hr.

**Figure 5.**
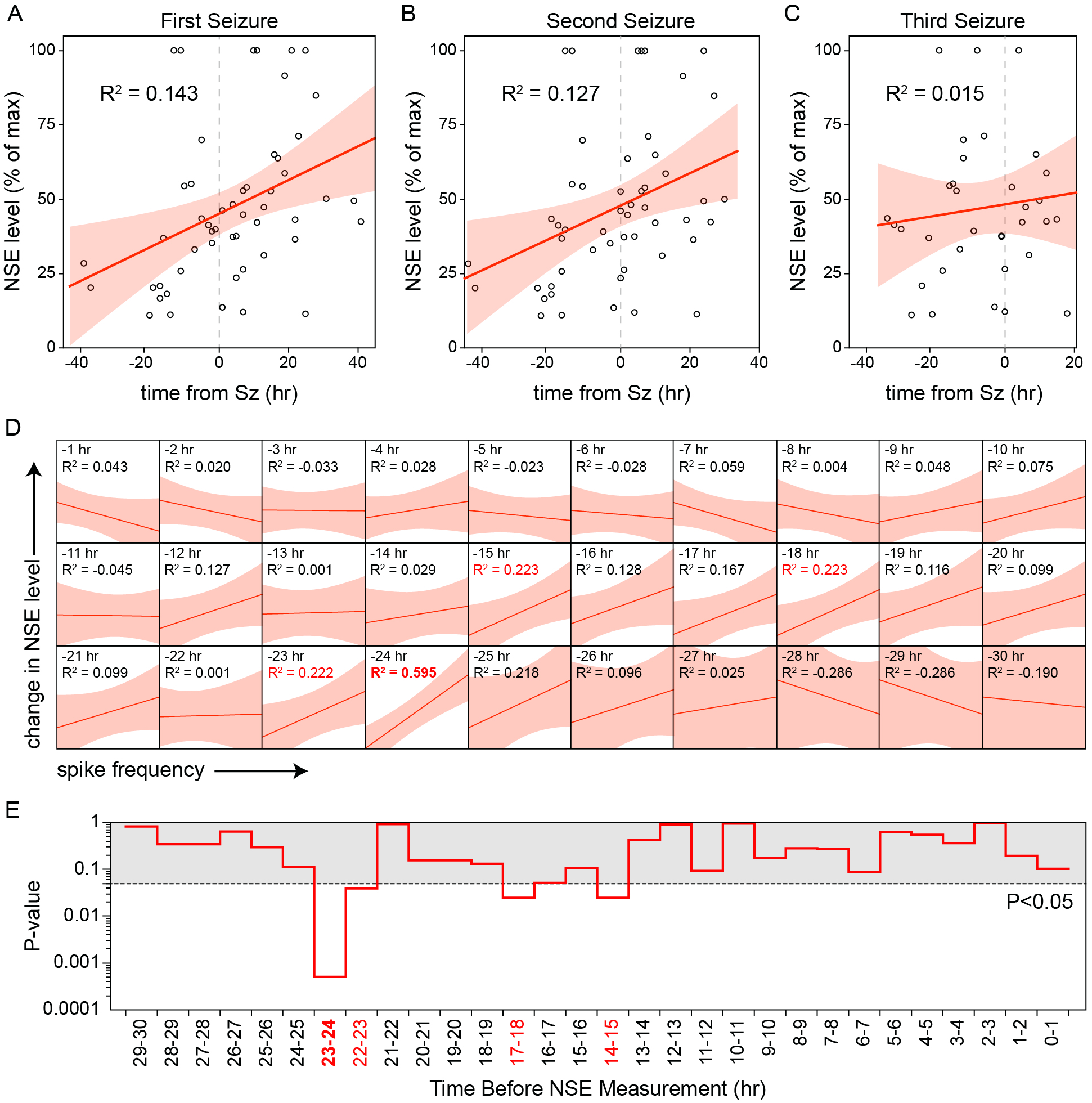
NSE levels increase after seizures and after increased spiking on the EEG. A) NSE measurements were converted to percent of maximum for each individual. The time of the first clinical seizure was set to t=0 and the normalized NSE measurements were plotted against number of hours before or after the seizure (each circle represents one NSE measurement). The distribution was analyzed by least squares linear regression to fit a line (red). The 95% CI for the regression is shown in solid light red. The fit (R^2^=0.143) indicates that NSE values were higher after the first seizure than before. B) The same process was applied to values relative to the second seizure time. As with the first seizure, NSE levels were higher after the second seizure than before (R^2^=0.127). C) The same process was applied to the the third seizure time. By the third seizure there was no longer a relationship between time after seizure and elevated NSE levels (R^2^=0.015). D) Continuous spike frequency data were available from EMU patients E003-E007. These data and the NSE measurements were recast as unitless values indicating increased serum concentrations or increased spike frequencies. Using standard least squares linear regression the NSE values were modeled relative to time of spike frequency measurement. Each panel shows the fitted line (red) and 95% CI for the regression (solid light red) from 0-1 hr (“-1 hr) before the serum measurement through to 29-30 hr (“-30 hr) before the serum measurement. E) The P-value derived from the *χ*^2^ analysis of a binomial maximum likelihood estimator is plotted against time relative to NSE measurement to reveal the most significant temporal epochs. The gray region shows P-values greater than 0.05; the dashed marks P=0.05.

## Discussion

Neuron-specific enolase, representing 1.5% of total soluble brain protein, is a ~78 kDa enzyme found predominantly in neurons and neuroendocrine cells^24; 25^. Enolases (2-phospho-D-glycerate hydrolases) are catabolic glycolytic enzymes that convert 2-phosphoglycerate to phosphoenolpyruvate as part of the cellular mechanism for ATP production^26^. Functional enzymes are formed by homo- and heterodimerization of α, β, and γ subunits differentially expressed in every cell type, with the neuron-specific form of enolase comprised of a γ-γ homodimer^26; 27^. Under normal conditions NSE levels in serum should be zero. However, ELISA-based methods for measuring NSE rely upon antibody recognition of the γ subunit, which is also found in platelets and erythrocytes, predominantly as an α-γ heterodimer^28^. As a result, baseline levels of γ-enolase in serum are approximately 10 ng/mL ^29^; in our study, healthy control values ranged from 3-22 ng/mL. During neurologic disease states, increased serum NSE is predictive of outcome and correlated to injury severity. For example, in closed-head traumatic brain injury (TBI), ~80 ng/mL NSE correlated with severe TBI, ~55 ng/mL correlated with moderate injury, and ~20 ng/mL was associated with mild head trauma^30^. Moreover, in this same study the level of serum NSE was 87% sensitive and 82% specific in predicting poor outcome. For the majority of trauma-related studies, including extracorporeal circulation-induced injury associated with cardiac surgery, the peak level of NSE was measured within 6-12 hours of the inciting event, slowly decaying with an apparent half-life of 24-48 hours^31^. This pattern suggests an accumulative building of NSE in the serum over the first few hours after injury followed by a gradual decline that is the sum of ongoing injury-dependent release and catabolic degradation of the enzyme in circulation. However, this pattern is at odds with our observations, in which large increases in NSE were detected within the space of 3 hours and large decreases occurred over similar time frames. Our findings suggest acute but transient neuronal injury events that result in a rapid spike of serum NSE followed by rapid decay of the existing NSE without ongoing replacement by continuous neuronal injury.

The assessment of NSE levels at multiple timepoints over the course of several days provided an unbiased dataset that upon post hoc analysis revealed a correlation between seizure and spike events and concomitant rises in serum NSE levels. By comparison to simultaneous measurement of S100β in the same subject along with similar temporal profiling in healthy control subjects, we identified statistically significant NSE signal changes in the epilepsy patients in our study. These findings are strengthened by the general stability of the S100β measurements through time, which rules out sample quality variability as an explanation for the NSE changes. Moreover, while all four control subjects exhibited signal variation values indicative of no change (1.0 or less), all 7 epilepsy patients had values above 1.0 (Figure 4B). Comparison of the 3 patients with low values (<1.5) versus the 4 patients with high values (>1.5) revealed no effect of age ([25 − 49 y] vs [31 − 45 y]) or disease duration ([5 mo − 29 y] vs [4 − 41 y]). The low variation in at least one patient (E007) is likely the result of an algorithmic false negative caused by the presence of two spikes in NSE level separated by a time window that masks the sample entropy difference (Figure 2H). Likewise, the low variation score in E003 may arise from the relative “noisiness” of the NSE measurements in this individual (Figure 2C), while the lower variation value measured in E002 may arise from the narrow dynamic range of the change in this patient (Figure 2B). Alternatively, these individuals may have different underlying etiologies or seizure foci/semiologies that preclude neuronal injury or there may be masking effects associated with different drug regimens or comorbidities. Overall, we are unable to determine whether all patients with temporal lobe epilepsy experience ongoing neuronal injury associated with seizures, but our findings support the presence of such injury in at least some patients.

Others have measured NSE and S100β in epilepsy patients, though none of these studies employed the same longitudinal profiling strategy in both patients and healthy controls. A study from Palmio and colleagues showed a statistically significant increase in both NSE and S100β at around 6 hours after a seizure and provided evidence that this change occurred in patients with temporal lobe epilepsy but not in individuals with extra-temporal lobe epilepsy^17^. While this supports our findings, it is notable that the change in NSE following seizures in this study was from 8.4 pg/mL to only 13.5 pg/mL, averaged across all of the patients with temporal lobe epilepsy, and the maximum NSE value measured in the study was around 22 pg/mL. In contrast, our averaged measurements ranged from 7.6 pg/mL to 35.0 pg/mL and the maximum NSE level we measured was 117 pg/mL. Whether this difference reflects aspects of the patient cohort, the unbiased sampling strategy employed in our study, or variations in sample processing is unknown. Nonetheless, the Palmio findings along with a number of other published studies^15; 32; 33^ support the contention that at least some patients with epilepsy experience ongoing neurodegeneration triggered by individual seizures. This concept is nicely reviewed by Pitkanen and Sutula^2^.

While this study was strengthened by the direct comparison of epilepsy patient measurements to repeated samples collected from healthy control subjects under similar conditions (e.g. intravenous line placement rather than repeated venipuncture, collection under in-patient-like conditions), a number of potential weaknesses require cautious interpretation of the findings. The most significant caveat regards the absence of overnight serum samples. This precludes continuous evaluation of the changes in NSE, especially in patients with clinical seizure events that occurred outside of the 6 AM to 6 PM collection window. Likewise, the absence of overnight serum samples may alter the correlation of spike frequency to NSE level. Obviously, these experiments are logistically quite challenging and expensive to perform. In addition to the demands on clinical personnel required for continuous sampling every 3 hours for up to 72 hours or more, the need to prepare each serum sample immediately after collection requires a concerted round-the-clock laboratory effort. In the absence of some sort of indwelling NSE sensor, however, all such studies will be limited by the sampling frequency and the difficulty of comparing a continuous measurement (EEG) to a discontinuous measurement (serum factors). In addition, another potential issue in this study was the collection of clinical quality EEG rather than research quality data. While we were able to perform automated spike frequency analysis in four of the seven EMU subjects, it is possible that the lower quality EEG restricted the sensitivity of the analysis. This suggests that future studies may benefit from either higher quality EEG, better algorithms for analysis of noisy EEG, or serum sampling in patients with intracranial electrodes. Finally, the methods employed for measuring NSE and S100β signal variation are challenged by the small number of samples and by sampling gaps. While our strategy for measuring sample entropy and signal variation accounts for the small sample size, this metric would benefit from more measurements and finer temporal resolution. A key example of the difficulties presented by a small sample size is the apparent false negative finding in E007, as discussed above. This patient exhibits a clear spike in NSE at the beginning of the study, but the second, albeit smaller, spike that occurs during the second day of measurements resulted in a low sample entropy score. Presumably, the availability of overnight serum samples would have filled the gap between these two spikes and enhanced the accuracy of entropy analysis. However, this problem at least suggests that the identification of high sample entropies and large signal variation metrics in the other patients are not false positives and were made despite a tendency of the algorithm and the gapped data to underestimate information content. The early NSE spike in patient E007 also reduced our ability to assess the impact of preceding seizures and EEG spiking events on changes in NSE levels, as we had less than 3 hours of EEG data collected before the NSE spike. Due to the post hoc nature of the serum analyses we were also unable to ascertain whether the subject had any relevant clinical seizures over the 24 hr preceding their enrollment into our study.

Despite these issues we did obtain several compelling associations. Increased levels of serum NSE were associated with increased time after the first seizure at P=0.0064. The coefficient of determination for this linear regression is 0.143, indicating that the relationship between time after seizure and increasing NSE levels is noisy. However, 100 iterations of 20% k-fold crossvalidation confirmed that this R^2^ value was significantly different from zero (95% confidence interval of the k-fold R^2^: 0.07 to 0.14; P<0.0001 by Wilcoxon signed rank test against a null hypothesis that R^2^=0; power = 0.999). Due to the discrete nature of both the seizure events and the serum measurements it is difficult to identify a specific post-ictal time domain for the increase in NSE. However, simple inspection of the plot in Figure 5A suggests that the NSE levels trend upward at around 20 hr after the first seizure. This time domain also appears to be relevant to the detection of increased NSE levels following increased spiking on the EEG. Visual inspection of Figure 5D suggests a broad, albeit low signficance, trend toward increased serum NSE from about 15 to 21 hr after an increase in spike frequency. Statistically, the strongest association between a preceding increase in EEG spiking and detection of increased serum NSE occurs at 24 hr. This time domain exhibited a strong coefficient of determination (R^2^=0.595), high statistical significance (P=0.0003), and high statistical power (0.9922), suggesting that despite the limitations of our current data we revealed a strong association between an electrophysiological disturbance and a concomitant rise in a neuronal injury marker in the serum after about 24 hr. Unfortunately, our ability to determine the length of time over which this rise in serum NSE persists after 24 hr is limited by the length of the EEG recording session for the EMU patients. Analysis of Figure 5D shows that by 27 hr after an increase in spike frequency our data are too sparse to draw interpretable conclusions (indicated by the broad 95% confidence interval bands (light red) around the regression fit (red line)). This suggests that future studies will need to retain the EMU subjects for longer EEG recording. This would also permit more serum measurements, further strengthening our ability to detect significant associations. Nonetheless, our current data support the strong, biologically relevant conclusion that an increase in serum levels of the neuronal injury marker NSE is detected approximately 24 hr after an electrophysiological event consistent with neuronal hyperactivity.

If our interpretation of these findings is correct, then post-ictal assessment of serum NSE may serve as a surrogate biomarker for measuring the efficacy of acute neuroprotective therapies aimed at preserving neurons in patients with epilepsy^34^. We recently reported that treatment of mice with an oral calpain inhibitor after the start of behavioral seizures induced by the neuroinflammatory response to acute viral infection resulted in preservation of hippocampal CA1 pyramidal neurons, preservation of cognitive performance, and abrogation of further seizure events^35^. Likewise, calpain inhibitor therapy started after onset of status epilepticus reduced seizure burden in the rat pilocarpine model^36^ and preserved CA1 neurons in the kainic acid model^37^. Because loss of hippocampal neurons, whether excitatory or inhibitory, may underlie the transition from spontaneous seizures to epilepsy as well as the persistence or spread of epileptic foci^34^, neuroprotective drugs may block epileptogenesis, prevent cognitive sequelae associated with seizures and epilepsy, and facilitate maintenance of seizure-free outcomes following brain resection surgery. However, directly measuring the efficacy of such neuroprotective drugs is challenged by time-to-effect and by the difficulty of correlating the absence of subsequent seizures, etc, to drug efficacy. We therefore propose that measurement of serum NSE will provide causal evidence of drug efficacy, particularly during acute post-ictal windows and perhaps especially in the context of a trial involving calpain inhibitor therapy delivered immediately after a seizure.

Finally, our findings provide further evidence of ongoing neuronal injury in patients with mesial temporal sclerosis, even in subjects with long disease durations. Because our study explicitly involved patients with intractable epilepsy, the measurement of seizure-associated NSE spikes in serum raises the question of whether neuron loss in these individuals is the *cause* of their intractable disease state. In other words, does the ongoing and accumulative low level injury of hippocampal neurons in these patients propagate neural circuit disruptions that render the system refractory to current drug strategies? If so, then initiation of neuroprotective therapy may effectively short-circuit a pathogenic feedback loop and convert even patients with long-standing intractable disease to a state that is amenable to standard treatment. Coupled with the obvious benefits for preventing cognitive decline, the potential to reverse intractability suggests that neuroprotective strategies must be more aggressively pursued in patients with temporal lobe epilepsy.

## Acknowledgements

Financial support was provided by grant NS064571 from the NIH/NINDS (CLH), grant UL1TR000135 from the NIH/NCATS, the Mayo Clinic Center for MS and Autoimmune Neurology, and a generous gift from the Albert and Mary Jane Staton Family. Materiel support was provided by the Mayo Clinic Center for Individualized Medicine. Expert technical assistance was provided by Sherry Klingerman, Jessica Sagen, Renee Johnson, Misha Patel, and Christina McCarthy.

## Author Contributions

S.K., R.G.L.-C., F.K., G.A.W, and C.L.H. collected and analyzed data. S.K. and C.L.H. prepared all figures; F.K. and C.L.H prepared the tables. S.K. and C.L.H wrote the manuscript. All authors reviewed the manuscript.

## References

1 Moshe SL, Perucca E, Ryvlin P, et al. Epilepsy: new advances. Lancet 2015;385:884–898.

2 Pitkanen A, Sutula TP. Is epilepsy a progressive disorder? Prospects for new therapeutic approaches in temporal-lobe epilepsy. Lancet Neurol 2002;1:173–181.

3 Breuer LE, Boon P, Bergmans JW, et al. Cognitive deterioration in adult epilepsy: Does accelerated cognitive ageing exist? Neurosci Biobehav Rev 2016;64:1–11.

4 Dingledine R, Varvel NH, Dudek FE. When and how do seizures kill neurons, and is cell death relevant to epileptogenesis? Adv Exp Med Biol 2014;813:109–122.

5 Cendes F, Sakamoto AC, Spreafico R, et al. Epilepsies associated with hippocampal sclerosis. Acta Neuropathol 2014;128:21–37.

6 Seidenberg M, Kelly KG, Parrish J, et al. Ipsilateral and contralateral MRI volumetric abnormalities in chronic unilateral temporal lobe epilepsy and their clinical correlates. Epilepsia 2005;46:420–430.

7 Kalviainen R, Salmenpera T, Partanen K, et al. Recurrent seizures may cause hippocampal damage in temporal lobe epilepsy. Neurology 1998;50:1377–1382.

8 Fuerst D, Shah J, Shah A, et al. Hippocampal sclerosis is a progressive disorder: a longitudinal volumetric MRI study. Ann Neurol 2003;53:413–416.

9 Caciagli L, Bernhardt BC, Hong SJ, et al. Functional network alterations and their structural substrate in drug-resistant epilepsy. Front Neurosci 2014;8:411.

10 Briellmann RS, Berkovic SF, Syngeniotis A, et al. Seizure-associated hippocampal volume loss: a longitudinal magnetic resonance study of temporal lobe epilepsy. Ann Neurol 2002;51:641–644.

11 Nairismagi J, Grohn OH, Kettunen MI, et al. Progression of brain damage after status epilepticus and its association with epileptogenesis: a quantitative MRI study in a rat model of temporal lobe epilepsy. Epilepsia 2004;45:1024–1034.

12 Lopim GM, Vannucci Campos D, Gomes da Silva S, et al. Relationship between seizure frequency and number of neuronal and non-neuronal cells in the hippocampus throughout the life of rats with epilepsy. Brain Res 2016;1634:179–186.

13 Kawata K, Liu CY, Merkel SF, et al. Blood biomarkers for brain injury: What are we measuring? Neurosci Biobehav Rev 2016;68:460–473.

14 Tanabe T, Suzuki S, Hara K, et al. Cerebrospinal fluid and serum neuron-specific enolase levels after febrile seizures. Epilepsia 2001;42:504–507.

15 Rabinowicz AL, Correale J, Boutros RB, et al. Neuron-specific enolase is increased after single seizures during inpatient video/EEG monitoring. Epilepsia 1996;37:122–125.

16 DeGiorgio CM, Gott PS, Rabinowicz AL, et al. Neuron-specific enolase, a marker of acute neuronal injury, is increased in complex partial status epilepticus. Epilepsia 1996;37:606–609.

17 Palmio J, Keranen T, Alapirtti T, et al. Elevated serum neuron-specific enolase in patients with temporal lobe epilepsy: a video-EEG study. Epilepsy Res 2008;81:155–160.

18 Fairbanks VF, Ziesmer SC, O’Brien PC. Methods for measuring plasma hemoglobin in micromolar concentration compared. Clin Chem 1992;38:132–140.

19 Scheuer ML, Bagic A, Wilson SB. Spike detection: Inter-reader agreement and a statistical Turing test on a large data set. Clin Neurophysiol 2017;128:243–250.

20 Richman JS, Moorman JR. Physiological time-series analysis using approximate entropy and sample entropy. Am J Physiol Heart Circ Physiol 2000;278:h2039–2049.

21 Pincus SM. Approximate entropy as a measure of system complexity. Proc Natl Acad Sci U S A 1991;88:2297–2301.

22 Yentes JM, Hunt N, Schmid KK, et al. The appropriate use of approximate entropy and sample entropy with short data sets. Ann Biomed Eng 2013;41:349–365.

23 Curran-Everett D, Benos DJ. Guidelines for reporting statistics in journals published by the American Physiological Society: the sequel. Adv Physiol Educ 2007;31:295–298.

24 Marangos PJ, Schmechel D, Parma AM, et al. Measurement of neuron-specific (NSE) and non-neuronal (NNE) isoenzymes of enolase in rat, monkey and human nervous tissue. J Neurochem 1979;33:319–329.

25 Kato K, Suzuki F, Umeda Y. Highly sensitive immunoassays for three forms of rat brain enolase. J Neurochem 1981;36:793–797.

26 McAleese SM, Dunbar B, Fothergill JE, et al. Complete amino acid sequence of the neurone-specific gamma isozyme of enolase (NSE) from human brain and comparison with the non-neuronal alpha form (NNE). Eur J Biochem 1988;178:413–417.

27 Rider CC, Taylor CB. Enolase isoenzymes. II. Hybridization studies, developmental and phylogenetic aspects. Biochim Biophys Acta 1975;405:175–187.

28 Marangos PJ, Campbell IC, Schmechel De, et al. Blood platelets contain a neuron-specific enolase subunit. J Neurochem 1980;34:1254–1258.

29 Casmiro M, Maitan S, De Pasquale F, et al. Cerebrospinal fluid and serum neuron-specific enolase concentrations in a normal population. Eur J Neurol 2005;12:369–374.

30 Meric E, Gunduz A, Turedi S, et al. The prognostic value of neuron-specific enolase in head trauma patients. J Emerg Med 2010;38:297–301.

31 Johnsson P, Blomquist S, Luhrs C, et al. Neuron-specific enolase increases in plasma during and immediately after extracorporeal circulation. Ann Thorac Surg 2000;69:750–754.

32 Willert C, Spitzer C, Kusserow S, et al. Serum neuron-specific enolase, prolactin, and creatine kinase after epileptic and psychogenic non-epileptic seizures. Acta Neurol Scand 2004;109:318–323.

33 Tumani H, Otto M, Gefeller O, et al. Kinetics of serum neuron-specific enolase and prolactin in patients after single epileptic seizures. Epilepsia 1999;40:713–718.

34 Loscher W, Brandt C. Prevention or modification of epileptogenesis after brain insults: experimental approaches and translational research. Pharmacol Rev 2010;62:668–700.

35 Howe CL, LaFrance-Corey RG, Mirchia K, et al. Neuroprotection mediated by inhibition of calpain during acute viral encephalitis. Sci Rep 2016;6:28699.

36 Lam PM, Carlsen J, Gonzalez MI. A calpain inhibitor ameliorates seizure burden in an experimental model of temporal lobe epilepsy. Neurobiol Dis 2017;102:1–10.

37 Araujo IM, Gil JM, Carreira BP, et al. Calpain activation is involved in early caspase-independent neurodegeneration in the hippocampus following status epilepticus. J Neurochem 2008;105:666–676.

